# Revisiting use of DNA characters in taxonomy with MolD - a tree independent algorithm to retrieve diagnostic nucleotide characters from monolocus datasets

**DOI:** 10.1101/838151

**Authors:** Alexander Fedosov, Guillaume Achaz, Nicolas Puillandre

**Affiliations:** A.N. Severtsov Institute of Ecology and Evolution, Russian Academy of Sciences, Leninsky prospect 33, 119071 Moscow, Russian Federation; Institut Systématique Evolution Biodiversité (ISYEB), Muséum national d’Histoire naturelle, CNRS, Sorbonne Université, EPHE, 57 rue Cuvier, CP 26, 75005 Paris, France

**Keywords:** DNA-based diagnosis, DNA character, DNA barcoding, taxonomy, formal description of taxa

## Abstract

While DNA characters are increasingly used for phylogenetic inference, taxa delimitation and identification, their use for formal description of taxa (i.e. providing either a formal description or a diagnosis) remains scarce and inconsistent. The impediments are neither nomenclatural, nor conceptual, but rather methodological issues: lack of agreement of what DNA character should be provided, and lack of a suitable operational algorithm to identify such characters. Furthermore, the reluctance of using DNA data in taxonomy may also be due to the concerns of insufficient reliability of DNA characters as robustness of the DNA based diagnoses has never been thoroughly assessed. Removing these impediments will enhance integrity of systematics, and will enable efficient treatment of traditionally problematic cases, such as for example, cryptic species. We have developed a novel versatile and scalable algorithm **MolD** to recover diagnostic combinations of nucleotides (DNCs) for pre-defined groups of DNA sequences, corresponding to taxa. We applied MolD to four published monolocus datasets to examine 1) which type of DNA characters compilation allows for more robust diagnosis, and 2) how the robustness of DNA based diagnosis changes depending on the sampled fraction of taxons diversity. We demonstrate that the redundant DNCs, termed herein sDNCs, allow for higher robustness. Furthermore, we show that a reliable DNA-based diagnosis may be obtained when a rather small fraction of the entire data set is available. Based on our results we propose improvements to the existing practices of handling DNA data in taxonomic descriptions, and discuss a workflow of contemporary systematic study, where the integrative taxonomy part precedes the proposition of a DNA based diagnosis and the diagnosis itself can be efficiently used as a DNA barcode. Our analysis fills existing methodological gaps, thus setting stage for a wider use of the DNA data in taxa description.

## Introduction

The application of names on living organisms, associated to their description, is an essential procedure to communicate their identities to the community of scientists and stake holders, and thus to bring them to the scope of scientific knowledge. The Linnaean system, applied following the rules of the relevant nomenclatural codes, is the most widely used system to formally describe living organisms. For example, according to the International Code of Zoological Nomenclature (ICZN), “to be available, every new name published after 1930 must be accompanied by a description or definition that states in words characters purported to differentiate the taxon…”. And further: “When describing a new taxon, an author should make clear his or her purpose to differentiate the taxon by including with it a diagnosis, that is to say, a summary of the characters that differentiate the new nominal taxon from related or similar taxa.” (Article 13.1.3). Whereas traditionally taxa descriptions are mainly based on morphological data (Dunn et al. 2003; Cook et al. 2010), the non-morphological characters, such as characters of DNA, if referred to in words, are equally accepted by all nomenclatural codes (Cook et al. 2010, Renner, 2016). The amount of DNA sequence data available to taxonomists is steadily growing since two decades, and is being accumulated with ever-increasing rates with the recent advent of high throughput sequencing. Currently DNA sequence data are sometimes more accessible than rare taxonomic expertise (Cook et al. 2010), and it is thus not surprising that DNA data is now widely used in phylogenetics, species delimitation (Fujita et al. 2012; Puillandre et al. 2012; Pante et al. 2015a) and specimen identification (Herbert et al. 2003; Janzen et al. 2009; Goldstein & De Salle 2011). Conversely, the use of DNA data in formal descriptions remains scarce, with a difference of two orders of magnitude when comparing the number of species described with and without DNA data (Renner 2016). Indeed, even when new species are delimited with DNA characters, these characters are not systematically used in the corresponding species descriptions.

As a result of such inconsistency, new lineages revealed though phylogenetic and species delimitation approaches typically enter “taxonomic purgatory”: discovered and published, but not established taxonomically (Goldstein & DeSalle 2011; Pante et al. 2015b). This persistent backlog in taxonomic treatments results from various reasons, such as the lack of support for the new taxa (Pante et al 2015b), publication strategy that will lead authors to postpone the descriptions (Agnarsson & Kuntner M 2007; Pante et al 2015b), unwillingness of molecular biologist to describe taxa (Pante et al 2015b, Satler et al 2013), or the deficiency of taxonomic knowledge, often referred to as ‘taxonomic impediment’, leading to a situation where specialists cannot cope with the increasing amount of data coming from relevant DNA-based studies (Taylor 1983; Goldstein & DeSalle 2011; Fontaine et al. 2012; Pante et al. 2015b). Finally, a frequently evoked reason is the lack of morphological difference among lineages revealed through the examination of non-morphological (usually, DNA sequence) data (Schlick-Steiner et al. 2007; Pante et al. 2015b) – a reasoning that in fact is not justified by the nomenclatural codes. In all these cases, when new taxa molecularly delimited are not formally described, the use of DNA characters in addition to morphological characters would speed up and improve taxonomic descriptions. DNA taxonomy was criticized extensively in the past (Dunn 2003 and references therein; Moritz & Cicero 2004; Rubinoff & Holland 2005; Rubinoff et al. 2006), however, use of DNA data for taxonomic description **in concert** with integrative approach to taxa discovery, would not replace, or create a parallel to the morphology based taxonomy (Will et al. 2005; Jörger & Schrödl. 2013), but will improve quality and usability of the taxonomic descriptions. In this perspective, some recent publications strongly recommend promoting DNA-based diagnoses by relevant taxonomic codes, which would allow same lines of evidence to be used for both, differentiation of a taxon from relatives, and its formal description, thus enhancing integrity of systematics in the ‘molecular era’ (Tautz et al. 2003; Goldstein & DeSalle 2011; Renner 2016).

Essentially, there are two related methodological gaps: Lack of a generally accepted practice in presenting DNA characters in diagnoses (i.e. what kind of characters, and how they should be presented), and the lack of the operational algorithm to compile DNA based diagnoses. Published studies presented three major types of DNA-based diagnoses: i) genetic distances to closest relatives, ii) diagnostic stretches of DNA sequences, and iii) specific substitutions (Goldstein & DeSalle 2011). None of them is methodologically flawless. Genetic distance, is relative value, and does not provide discrete traits that can be associated with a diagnosed taxon (ideally to its type material). The stretches of DNA sequence inevitably contain uninformative nucleotide positions together with informative ones, and thus do not strictly qualify as diagnoses, but more as descriptions. Thus, providing specific substitutions is probably best practice of DNA character presentation in the diagnosis. Nevertheless, the term ‘substitution’ has implicit comparative context, but the framework of this comparison is seldom defined clearly and/or limited to one or few already described taxa (Zielske & Haase 2015), making the diagnosis meaningless in a broader context. The lack of compatibility between the three types of DNA characters presentation is hampering their consistent use in taxonomy, it is therefore important to establish common practice of providing DNA based diagnosis with its scope and mode of application explicitly defined. Furthermore, most published studies (Jörger & Schrödl 2013; Churchill et al. 2014 and Zielske & Haase 2015), report pure single nucleotide characters – i.e. nucleotides at specific positions in the DNA alignment, each being shared by all members of a diagnosed taxon, and by none of the other taxa in the data set. This approach allows for concise and robust diagnoses, but can only be applied to data sets of few species. In the data sets comprising hundreds of species recovery of even one such position for each species is unlikely. So, either, more taxonomically restricted data sets should be considered, or one should opt for use of composite characters (i.e. set of several specific nucleotides that together comprise a diagnostic character – see Material and Methods). The latter approach would be preferable in the highly diversified, poorly studied or taxonomically problematic groups – i.e. in the cases where defining a restricted scope of analysis can be difficult. Currently only one computational tool, CAOS (Sarkar et al. 2008) is capable of recovering composite DNA characters, but implemented through a tree based algorithm, it has some inherent limitations (see below), and is difficult to run. We developed a versatile and efficient algorithm MolD (for MOLecular Diagnosis) to identify diagnostic nucleotide characters for *predefined groups* of DNA sequences in large monolocus data sets. Predefined group of sequences in this context represents a species or a higher rank taxon, which we seek to diagnose. MolD is a tree independent algorithm (unlike CAOS), and therefore it i) allows to diagnose taxa that are not recovered as monophyletic in monolocus dataset (see below), and ii) does not require manual adjustments to run. In turn, crucial advantage of MolD over nucDiag in the SPIDER package for R (Brown et al. 2012) is that MolD identifies composite diagnostic characters (DNCs), which is crucial for larger data sets where no pure characters can be identified (see Discussion).

We applied MolD to several published DNA data sets aiming to assess usefulness of the routinely sequenced barcode DNA markers to supply diagnostic DNA characters for taxa of different ranks and to evaluate two possible types of DNA based diagnoses in terms of operationability for taxonomic purposes and robustness.

A diagnosis retrieved after the analysis of a finite data set is only *assumed* to be valid descriptors of the respective taxa in general. The validity of this extrapolation depends on how accurately the initial data set conveys diversity of the studied group. So, the third questions that we address is how the sampling coverage affects robustness of DNA-based diagnoses, and whether universal minimal requirements for the taxonomic sampling can be formulated to ensure sufficiently reliable diagnoses.

## Material and Methods

### Term definition

The taxon being diagnosed from here onwards is termed **focus taxon**, and is opposed to all other taxa in the dataset that are referred to as **non-focus taxa**. Single nucleotide characters can be classified into **pure diagnostic sites** (Sarkar et al. 2008), i.e. positions in the nucleotide alignment in which all members of a focus taxon share a given nucleotide, and all members of non-focus taxa have different nucleotide(s). On the figure 1 position 256 is a pure diagnostic site for the focus taxon *Conasprella.* Unlike pure diagnostic sites, the **private** characters (Sarkar et al. 2008) are the nucleotide positions at which some (but not all) members of the focus taxon, but none of the non-focus taxa members, share a given nucleotide (Fig 1, positions marked by asterisks). In addition, we introduce a new category of characters that we term **nested** characters, i.e. sites at which focus taxon members and some (but not all) non-focus taxa members, share a given nucleotide (positions in Fig. 1 marked with grey arrow). A **pure composite** nucleotide character – i.e. a combination of nucleotides at specific positions in the DNA alignment that are shared by all members of a focus taxon, and by none of the non-focus taxa members, is termed **primary Diagnostic Nucleotide Combination**, or **pDNC** (each of the nucleotide combinations 1-4 highlighted on the fig 1 is a pDNC). A pDNC comprising one nucleotide position (combination 1 on the fig 1) corresponds to a pure diagnostic site. A pDNC can be viewed as a minimal sufficient condition to assign a DNA sequence to a focus taxon. As each of such characters is sufficient to differentiate a query taxon in the context of the analysed data set, each is equivalent of a complete diagnosis. Two pDNCs may overlap by one or several nucleotide positions (e.g., pDNCs 2 and 3 on the figure 1), or share no positions (e.g. pDNCs 2 and 4); in the latter case the two pDNCs are termed **independent** pDNCs. In the case that all identified pDNCs share one or more nucleotide positions (i.e. no independent combinations are identified), such position(s) present in all pDNCs are termed **key positions**. Key position(s) appear(s) crucial for diagnosing a taxon, because a substitution at this position even in one sequence attributed to a focus-taxon immediately makes the focus-taxon impossible-to-diagnose with the selected genetic marker. On the contrary, when *n* independent pDNCs are recovered, no less than *n* substitutions would be needed to make a focus taxon undiagnosable; the likelihood of the latter scenario is obviously much lower. It is reasonable to expect, therefore, that the robustness of molecular diagnosis is higher when several independent pDNCs are recovered and the number of independent pDNCs can be viewed as a proxy for the level of confidence of an inferred pDNC-based diagnosis.

**Figure 1.**
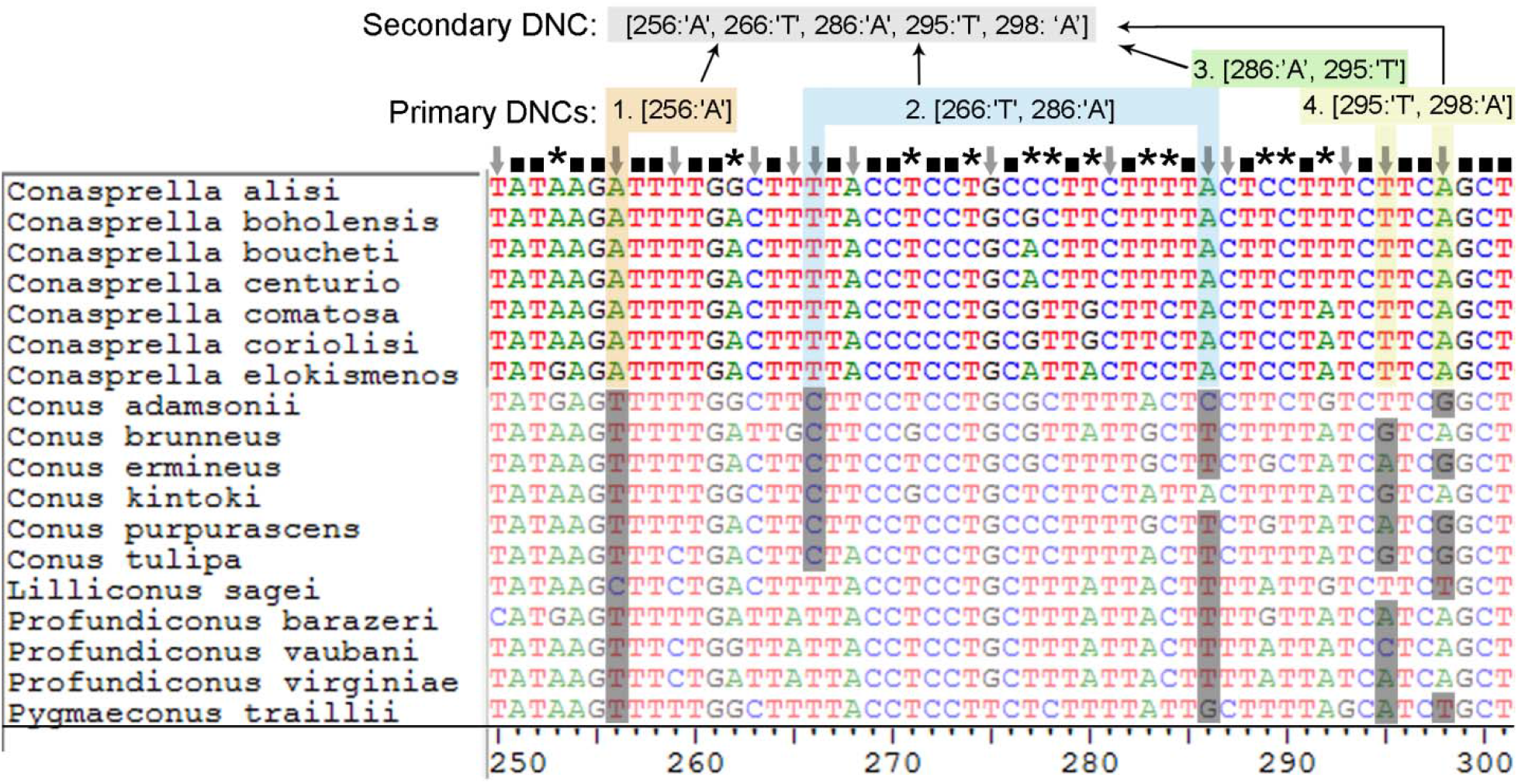
Major types of the DNA characters in the Conidae alignment; focus taxon *Conasprella.* Invariable nucleotide positions marked with black squares, nested and pure – with grey arrows, private – with asterisks. Nucleotide combinations 1, 2, 3, 4 (pDNCs) highlighted with colour; sDNC – with grey.

A diagnostic nucleotide combination, which **combines several pDNCs** (or characters from several pDNCs) aiming for an increased robustness of a diagnosis, is here termed **secondary DNC (sDNC)**. A sDNC compiled from the shown pDNCs is highlighted in the fig 1 in grey. As a whole, it fulfills the two main criteria of a DNC, 1) it is shared by all members of a focus taxon, and 2) it appears in none of the non-focus taxa members. However, unlike pDNCs, a sDNC is redundant and so the status of each nucleotide position in sDNC is not defined. The term ***diagnosis*** is used from here onwards to refer to either the multitude of the pDNCs, or to a single sDNC – a subset of informative nucleotide positions selected to convey the molecular identity of a taxon.

Most published studies (Jörger & Schrödl 2013; Churchill et al. 2014 and Zielske & Haase 2015), report a set of pure single nucleotide characters (i.e. a pDNC-based diagnosis, where each pDNC comprises one position only). We are not aware of any study providing composite DNA characters, either pDNCs comprising two or more nucleotide positions, or sDNCs. Furthermore, Jörger & Schrödl (2013) recommend abstaining from use of the composite characters, as the nucleotides that constitute them may represent contrasting substitution patterns. In the present study we examine in detail pros and cons of using pDNCs and sDNCs as DNA based diagnoses, and came to the conclusion that the sDNCs allow for more robustness, and therefore would be a preferable type of a DNA based diagnosis.

### Design of data analysis

The program MolD (**Mol**ecular **D**iagnoses) written in Python 2.7 is designed to retrieve pDNCs and sDNCs for the pre-defined assemblages of DNA sequences that correspond to taxa (which may range from species, subspecies, or even populations to any higher level). It constitutes the functional core and the main tool used in all analyses in the present study, combined with varying options of input and output.

Complete published DNA sequence data sets, partial data sets derived from the complete ones after random resampling of DNA sequences, or data sets of modified DNA sequences from original data were used as inputs. The sets of independent pDNCs, and 25 shortest pDNCs recovered for taxa of different rank (i.e. pDNC-based diagnoses), or redundant sets of DNA characters (sDNC-based diagnoses) were the two main types of output. Depending on what each particular analysis was aimed at, we also recorded proportion of variable sites in the DNA alignment for the focus taxa, validity of a diagnosis recovered for a partial data set against the respective complete data set. All analyses were implemented in Python 2.7 scripts, and run in Spider environment for Python. All scripts used for the present study are available in the supplementary data. The command line based version of MolD can be used as is, and it is currently being developed into graphical interface enhanced software.

### Data sets

Both the protein-coding and non-coding genes can potentially be used to retrieve DNA-based diagnoses (Jörger & Schödl 2013). Nevertheless, aligning sequences of non-protein coding genes unambiguously is labor-intensive (see Jörger & Schödl 2013) and not always robust, especially if data sets comprise highly divergent lineages. As the subjective component, which is unavoidable when aligning non-protein coding sequences, may notably affect the output diagnostic nucleotide characters, we avoid using them, and only consider relatively quickly evolving protein-coding genes – i.e. those typically used as DNA-barcodes in animals. These markers are commonly sequenced for large assemblages of specimens, and therefore, vast data sets can be accessed in publicly available data repositories.

We analysed three published data sets (**main data sets**) from highly divergent taxa: the dragonflies and damselflies (Odonata) (Bergmann et al. 2013), the bird family Furnariidae (Derryberry et al. 2011), and the cone-snails (Mollusca: Gastropoda: Conidae) (Puillandre et al. 2014) (Table 1). Odonata exemplifies an ancient and highly diversified insect taxon, which has become a well-established model group for DNA barcoding studies (Rach et al. 2008; Bergmann et al. 2013, Rach et al. 2017; Koroiva et al. 2017). The family Furnariidae represents one of the best molecularly sampled bird radiations and thus constitutes a good model for assessing robustness of DNA-based diagnoses derived from partial taxonomic coverage. The family Conidae includes the relatively young genus *Conus,* which encompasses over 750 extant species and exemplifies a hyperdiverse taxon of marine invertebrates (Olivera et al. 2014; Uribe et al. 2017). The three main data sets differ drastically in the taxonomic coverage, from less than one percent of over 8,000 known dragonfly and damselfly species, to ∼97% of the 293 accepted Furnariidae species, with cone snails data set being intermediate (Table 1). Although three protein coding genes: COI, ND2, and ND3 were available for Furnariidae (Derryberry et al. 2011), and all three were tested, all presented analyses were performed on the ND2 data set, which contained longest sequences (1041 bp) and was the most informative. Both the ND1 and CO1 were analysed for Odonata (Bergmann et al. 2013), and only CO1 sequences were available for Conidae.

**Table 1.** Analysed data sets.

| Dataset | Marker<br>(n.p.) | Main data set<br>N spp / N seq | Reference | Species<br>coverage | Expanded data set<br>N spp / N seq |
| --- | --- | --- | --- | --- | --- |
| Odonata<br>(Order) | COI (541) | 46 / 234 | Bergmann et al. 2013 | <1% | 816 / 4029<br>(GenBank) |
|  | ND1 (316) | 50 / 266 | Bergmann et al. 2013 | <1% | --- |
| Furnariidae<br>(Family) | ND2 (1041) | 275 / 293 | Derryberry et al. 2011 | 97% | 285 / 788<br>(GenBank) |
| Conidae<br>(Family) | COI (658) | 230 / 1438 | Puillandre et al. 2014 | 25% | 230 / 1894<br>(Duda et al. 2012) |

While most analyses were performed on these main data sets, we retrieved additionally multiple COI sequences of the 4 species of *Conus* for which a population structure analysis was performed (Duda et al. 2012), as well as all ND2 sequences of Furnariidae, and most COI sequences of Odonata available in the GenBank. Thus complemented **expanded** data sets **Furnariidae_788ND2**, **Odonata_4kCOI**, and **Conidae_COI_4spp** were built (Table 1). The three expanded data sets were used to assess the reliability of the diagnoses recovered on the smaller main data sets. Up to 15 ambiguously called nucleotides were allowed in the Furnariidae_788ND2 and Odonata_4kCOI data sets, and up to 25 in the Conidae_COI_4spp. The criterion of the sequence quality was relaxed deliberately compared to the main data sets to test the robustness of the recovered DNA based diagnoses in the data sets including some low-quality DNA sequences.

As all DNA sequence data sets used here represent protein coding genes, the alignment (performed by Clustal ω at https://www.ebi.ac.uk/Tools/msa/clustalo/) was straightforward, and could not potentially become a source of incorrect homology hypotheses. Subsequently, a RaxML tree with 1,000 bootstrap iterations (Stamatakis 2006; Miller et al. 2010) was constructed for each data set and used to check that the sequence attributions to taxa are consistent across the data set. As was said above, a focus taxon might appear paraphyletic, which would not make it *a priori* undiagnosable, but attribution of identical sequences to two or more different taxa, resulting, for example, from misidentification, will make these taxa undiagnosable. All outgroup sequences were removed before the application of MolD.

On the first stage we applied MolD to the three main data sets to characterize the output pDNC-based diagnoses for taxa of the different rank: a) All species of Odonata in both the COI and ND1 data sets; b) Odonata families Aeshnidae and Libellulidae in both the COI and ND1 data sets; c) Furnariidae genera *Synallaxis* and *Cranioleuca;* d) Furnariidae subfamilies Sclerurinae, Dendrocollaptinae and Furnariinae; e) Five species of Conidae represented in the main Conidae data set by 19 or more sequences; f) Subgenera of *Conus: Cylinder, Pionoconus, Rhizoconus* and *Turriconus;* g) Genera of Conidae: *Conus* and *Conasprella.* When the diagnoses were recovered, three output metrics were recorded: total number of the recovered pDNCs, number of independent pDNCs in the diagnosis (Nind), and number of nucleotide positions in the shortest pDNC.

### The effect of taxonomic sampling on the reliability of the DNA based diagnosis: expectations

We used random taxon resampling on published DNA datasets to assess the robustness of DNA-based diagnoses associated to different sampling fractions. Our rationale was that if we access a sufficiently large DNA sequence data set of some taxon, this data set may be used to model finite genetic diversity of this taxon. Then, we can select few focus taxa, representing both stem and crown group lineages of this taxon, and recover their DNA-based diagnoses from the entire data set. These diagnoses, although only partly capture identity of the respective focus taxa, in the framework of the model can be viewed as *finite* diagnoses. Then, by providing diagnoses to subsamples of this large data set, and comparing each of them against the finite diagnosis, the robustness of a diagnosis associated with each sub-sample can be estimated. Using this approach we were sampling increasing fractions of four published data sets, each – for several focus taxa. The sub-samples of the entire dataset from here onwards are referred to as *partial* datasets. The robustness of the diagnosis compiled from a partial dataset is measured by the proportion of the entire-data-set-valid pDNCs, *EDV-pDNCs,* in this diagnosis – i.e. those pDNCs that although retrieved from a partial data set, remain valid pDNCs in the context of the entire data set. We expect that this proportion will be very low when few sequences or taxa are sampled, but as more sequences are added to the data set, the proportion of the EDV-pDNCs will increase until finally reaches 100% when all the diversity is sampled. The curve describing the proportion of the EDV-pDNCs as a function of the sampling fraction may reach a plateau earlier: in this case the sampling fraction at which the plateau is reached would mark the minimum taxonomic sampling sufficient to provide a robust DNA diagnosis. We expected thus that the resampling graphs will be similar to typical accumulation curves – the sampled genetic diversity of a focus taxon will grow quickly in the beginning and subsequently slower.

The taxon resampling was performed separately for the species level and supra-specific focus taxa (see two following sections). In the case of species we reformulated the question to ‘how many haplotypes need to be included to propose a credible diagnosis of the species’? If leaving population structure aside, for the supra-specific taxa, the sampling should ideally be representative of the diversity at two hierarchical levels, intraspecific and interspecific, and would translate into number of species included, and number of haplotypes in each species included. The data sets with multiple sequences (and thus with multiple haplotypes) per species will show higher level of the genetic diversity of both focus taxon and non-focus taxa, and so, would better approximate real genetic diversity of an analysed lineage. This would expectedly lead to the higher proportion of the EDV-pDNC at each sampling fraction, and so if the robustness threshold is to be reached, we expected it to be reached earlier in the case, when multiple sequences per species are included.

### pDNC-based diagnosis for species: haplotype resampling

We carried out iterated resampling for the unique COI haplotypes of the *C. chaldaeus, C. ebraeus, C. miliaris* and *C. sanguinolentus* in the Conidae_COI4spp data set. At each iteration, *n* unique COI haplotypes were picked up randomly for a focus species, and thus created partial data set was completed by all records of Conidae_COI4spp for non-focus taxa. Thus obtained data set was passed to MolD to retrieve a set of pDNCs. The number of sampled unique haplotypes *n* was increasing from 2 and up to the total number of unique haplotypes available for each species, and was incremented by one until *n* equals 20, and by five subsequently. Ten iterations have been made for each tested number of unique haplotypes. The output parameters recorded after each iteration were 1) number of the COI sequences of the focus species sampled, 2) the proportion of variable sites among the sampled sequences of the focus taxon (hereafter PVS), as a measure of the molecular diversity of the sampled species, 3) the number of independent diagnostic nucleotide combinations sampled (NInd), and 4) the total number of pDNCs retrieved. Each of the pDNCs retrieved for the partial data set, was checked against the full data set to identify whether it is an EDV-pDNC. This test allowed estimation of how the reliability of the pDNC-based diagnoses compiled for the subset of sequences changes when an increasing proportion of the genetic diversity of species is sampled. However, the results of this analysis are only valid under the assumption that the final data set available encloses the entire genetic diversity of the group (for discussion see Wiens & Servedio 2000 and Rach et al. 2008).

### pDNC-based diagnosis for supraspecific taxa: species resampling

We performed iterated taxonomic resampling for the subfamily Sclerurinae and the genus *Synallaxis* (Furnariidae) and for the genera *Conus* and *Conasprella* (Conidae). The expanded data set Furnariidae_788_ND2 was used for Sclerurinae and *Synallaxis* to enable inclusion of multiple sequences per species, and the main Conidae data set was used for *Conasprella* and *Conus.* An increasing number of species of the diagnosed taxon has been picked up randomly, starting from 2 and up to the total number of species in the data set, and a partial data set was thus built from the sequences of the picked species. Similar to the haplotype resampling, for each tested number of species, 10 iterations have been performed, and the increment was set to 1 for the Sclerurinae, *Synallaxis* and *Conasprella* (2, 3, 4 species resampled, and so on), or to 5 for *Conus* (2, 7, 12…). The resampling was conducted under two different regimes in respect to the focus taxon – either all sequences of a picked species were added to the partial data set, or each picked species was represented by single sequence. By comparing the outputs of the resampling under these two regimes we were able to estimate how the number of sequences per species in the input data affects reliability of the resulting diagnosis. The number of species sampled from each of the non-focus taxa was increasing with different increment, so that the proportion of species sampled from each taxon remained equal. In each iteration, a MolD run was performed on the sampled partial data set to retrieve a set of pDNCs for the focus taxon. The output parameters recorded for each iteration were 1) number of species of the focus taxon sampled, 2) total number of species sampled, 3) PVS in the focus taxon sequences, 4) total number of the pDNCs sampled, 5) Nind, 6) Proportion of the EDV-pDNCs.

### Assessment of robustness of sDNC-based diagnoses

Since taxonomic sampling has a profound effect on the reliability of the DNA based diagnosis, another crucial question is how a growing genetic diversity associated with increasing taxonomic sampling can be simulated without increasing the actual number of sequences in the data set. Such algorithm would allow to test and refine the DNA-based diagnoses having only a limited number of specimens actually sequenced. We utilized the approach described by Bergmann and co-workers (2013) and generated artificial DNA sequence data sets to test the robustness of recovered DNA based diagnoses. We were introducing a pre-defined number of random nucleotide substitutions at random sites of a nucleotide sequence, without phylogenetic pattern, and not at the sites recognized as key positions. By doing so, we generated artificial data sets that are partly or entirely composed of the modified sequences of our main data sets (for parameters used for species-level and supraspecific taxa, see below). By generating 100 random artificial data sets and checking whether a DNC retrieved for a focus taxon from the actual sequence data stands in the context of each artificial data set, we were able to evaluate the robustness of the DNC. Here we allowed 1 mismatch in the positions involved in the sDNC in a focus taxon sequences: in this case the sDNC was still considered valid, if remaining positions constituted a valid DNC. Each sDNC was thus scored by assigning a number from 0 (when a DNC failed in all 100 artificial data sets) to 100 (when it worked for all 100).

The explained algorithm to score robustness of DNCs was used to transform a set of pDNCs into single sDNC. Unlike pDNCs, sDNCs are redundant (see terms), and therefore are expected to constitute more robust diagnoses. To transform a pDNC-based diagnosis into an sDNC, the remaining informative nucleotide positions in the alignment were ranged by how frequently they occur in the DNCs. Then, either a list of single-position pDNCs (if exist), or the shortest retrieved pDNC was taken, and expanded step-wise by appending most frequently recovered nucleotide positions. Each time a new position was added, the resulting sDNC was scored using 100 artificial data sets, and either send to output (if the sDNC meets pre-defined robustness criteria detailed below) or another position is appended to it, and so on until the robustness criteria are met. When all informative nucleotide positions are appended to the sDNC, or the length of sDNC exceeds the arbitrarily set limit of 25 positions, and the required confidence threshold has not been reached, the sDNC is output with a warning.

### sDNCs in diagnoses of species level taxa

The use of artificially generated DNA data sets allowed us to model underrepresented genetic diversity in both the focus- and non-focus taxa. When addressing this issue at the species level, each sequence in the main data set was modified, and the number of substitutions was such that the derived artificial sequence would be 1% different from the original one. One percent difference is well within the typical K2P genetic distance for within species comparisons of the analysed taxa (Bergmann et al. 2013), and therefore, each artificial sequence would be confidently attributed to the same species. However, as the modifications of each sequence are random, this effect reproduced in 100 artificial data sets creates a more complex genetic diversity landscape to test robustness of the sDNCs. A sDNC was output when it has scored 99 or 100 in three consecutive tests, which may be a very strict setting, but worked efficiently in the analysed data sets. Then, to evaluate robustness of an output sDNCs (retrieved from the main data sets), it was validated against respective expanded data set in the same manner, as the validity of the pDNCs was tested (see above). This analysis was performed for 15 species of Odonata, which were represented in the main Odonata COI data set by six or more sequences, and for four species of *Conus,* genetic diversity of which was notably better represented in the Conidae_COI4spp data set. Thus, we estimated whether the entire approach (data sets of artificially generated sequences), and the particular parameters that we employed to rate the sDNC-based diagnoses, can efficiently model the missing genetic diversity in the data set. The output that meets the criteria of robustness would suggest that the effect of the partial taxonomic sampling is leveraged, and the resulting DNA-based diagnosis can be considered as robust.

### sDNCs in diagnoses of supraspecific taxa

For supraspecific taxa the number of random substitutions was set to make each of the modified sequences 3% different from the original one. Such level of the divergence applied to each sequence in the original data set has led to overlaps in molecular identities of closely related taxa, and no sDNC could pass any reasonably high threshold. Therefore, only 10 percent of the sequences in each main data set, and in no more than 20 species per taxon (numbers selected arbitrarily) were modified. A sDNC was output when it had scored more than 85 in three consecutive tests.

Finally, we coupled the described algorithm of stepwise evaluation of the sDNC robustness with taxonomic resampling algorithm explained above to estimate how the credibility of the sDNC-based diagnoses changes with increasing taxonomic sampling. This analysis also enabled direct comparison with the resampling results obtained for the pDNCs. The same regime with all sequences of a picked species added to the partial data set has been used. The increment was set to 1 for Sclerurinae (1,2,3… until all the 17 species are sampled) and *Profundiconus* (Conidae), to 2 in *Conasprella* and *Synallaxis* (2,4,6…), and to 2 (in the range of 20) and subsequently changing to 5 in *Conus* (2,4,6,8,10,12,14,16,18,20,25,30, …185). Taxonomic resampling for cone snail taxa was performed on the main data set, and for Sclerurinae and *Synallaxis* – on the extended data set Furnariidae_788_ND2, to take into account multiple sequences per species. Ten iterations have been made for each sampled number of species. Validity of each of the 10 retrieved sDNC was checked against a complete data set, and it was assigned a value of 1 if it was an EDV-sDNC, or 0, if it was not. A mean value for 10 iterations was plotted as a function of the number of species sampled. Finlly the differences among the randomly generated artificial datasets that are used to rate sDNCs, will expectedly lead to certain degree of inter-run variation. Thus to enable the interrun comparison, output of each of the ten iterations was also recorded.

## Results

### pDNC based diagnoses for taxa of different rank

#### Odonata

The analysis of ND1 data set has led to identification of 349 to 1454 (mean 666) unique pDNCs comprising no more than 7 positions for all but two Odonata species (Fig. 2, Supplementary table 1). The exceptions are the species in the poorly genetically differentiated pair *Pseudagrion niloticum* – *Pseudagrion acaciae:* no pDNC were identified for either of the two species. The number of independent pDNC recovered for each Odonata species ranges from 4 to 25 (mean 16). The shortest pDNC comprising 1 position (i.e. a pure diagnostic site) was detected in 29 species; the mode of pDNC length distribution (analysed for all species with 10 or more sequences in a data set) was 3 nucleotides (Supplementary table 1). Very similar results were obtained with the CO1 data set – mean number of the pDNC of 779, with the number of independent pDNC ranging from 2 to 33 (mean 20), and shortest pDNC in most species comprising one nucleotide position only (Fig. 3, Supplementary table 2). The species pair *Pseudagrion niloticum* – *Pseudagrion acaciae* could not be differentiated based on the COI data set as well, but when all sequences of these two species were attributed to a single taxon, it could be easily diagnosed in both the ND1 and the CO1 data sets. The families Aeshnidae (11 and 10 species in the ND1 and COI main data sets respectively) and Libellulidae (16 and 19 species in the ND1 and COI main data sets respectively) were also successfully diagnosed, and the statistics of the diagnoses were generally comparable to the pDNCs recovered for species of Odonata (Fig. 2, Supplementary table 3)

**Figure 2.**
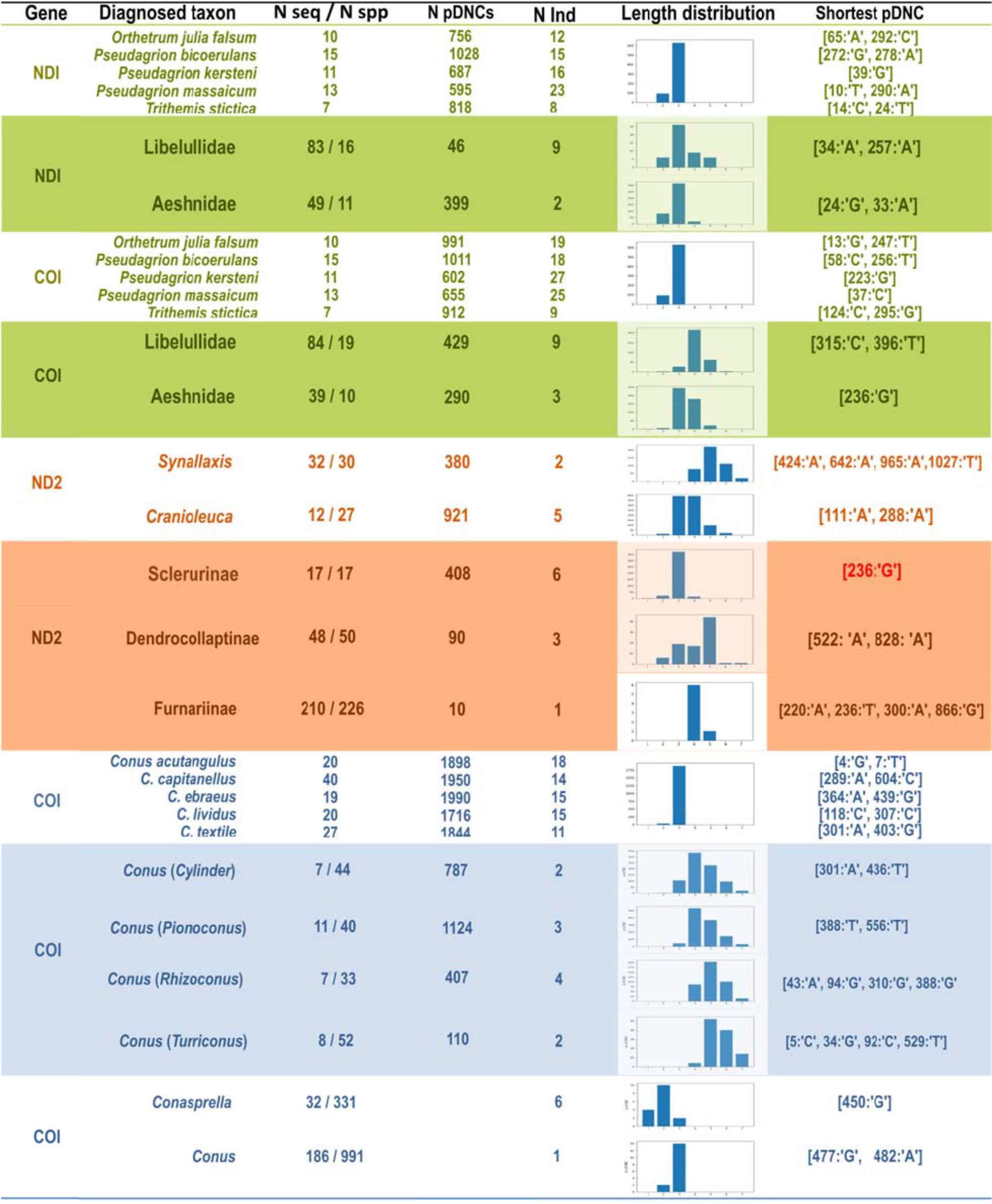
Metrics of pDNC-based diagnoses based on the analyses of main data sets, for species and supra-specific taxa.

#### Furnariidae

Diagnostic nucleotide combinations were successfully compiled for all five supraspecific taxa of Furnariidae. A total of 881 and 431 unique pDNCs were obtained for Cranioleuca and Synallaxis respectively (Fig. 2). The two genera differ notably in the number of recovered independent pDNCs (5 and 2 respectively), and the distribution of pDNC lengths in *Synallaxis* was notably shifted towards longer combinations compared to *Cranioleuca.* The number of pDNCs recovered for the subfamilies of Furnariidae ranged from 7 for Furnariinae (with a key nucleotide position 220: ‘A’) to 415 for Sclerurinae (6 independent). The shortest pDNC recovered for Sclerurinae in the main data set (17 Sclerurinae sequences) comprised one nucleotide position only, however it was not anymore a diagnostic combination of Sclerurinae in the expanded dataset (180 Sclerurinae sequences). No pDNCs shorter than 4 positions were retrieved for Furnariinae and *Synallaxis* in either main or expanded data set.

#### Conidae

The pDNC-based diagnoses compiled for five *Conus* species with 19 or more specimens from at least two geographic localities (Fig. 2, Supplementary table 4) were overall comparable with the diagnoses obtained for Odonata species, with the following differences: a) on average 14.6 independent pDNC were retrieved for a species of *Conus,* compared to over 20 in species of Odonata; b) no single-position pDNCs were retrieved for any of the diagnosed *Conus* species; c) the length distribution of the pDNC in *Conus capitanellus* and *C*. *textile* is clearly shifted towards longer combinations (Supplementary fig. 1). The lack of single-position diagnostic characters in *Conus* species is due to the notably larger number of species included in the data set (230 Conidae spp. VS 48-50 Odonata spp.). The number of informative sites for Odonata species varies from 165 to 194 in the ND1 alignment (i.e. 3.3 – 3.88 times as the number of species) and from 214 to 268 in the CO1 alignment (4.28 – 5.8 times the number of species). In the Conidae data set, the number of informative sites in the COI alignment (from 298 to 349) divided by to number of species (230) is thus notably smaller: 1.3 – 1.52.

Compared to species, fewer independent pDNCs were sampled for each of the four *Conus* subgenera: from 2 in *Turriconus* and *Cylinder* to 4 in *Pionoconus.* The pDNC length distribution is clearly shifted towards longer combinations (Fig. 2, supplementary table 5) with most pDNCs comprising 4 or 5 nucleotide positions. Finally, MolD has successfully diagnosed the conid genera *Profundiconus, Conasprella* and *Conus* (Fig. 2, supplementary table 6).

### The effect of taxonomic sampling on the robustness of the DNA based diagnosis

#### Robustness of the pDNC-based diagnosis for species level taxa

The 342 to 1289 pDNCs per species recovered for the 15 species of Odonata in the main Odonata COI data set were tested against the Odonata_4kCOI data set with almost 20 times larger number of species included (Supplementary table 8). The proportion of pDNCs that remain valid (i.e. shared by all focus taxon specimens) and diagnostic (i.e. appear in none of the non-focus taxa) in the expanded data set appears extremely low – from 0 to 3.7% (*Platycypha caligata*), mean 0.53%. In the case of Conus, the species lists of the main and expanded data sets were identical, but the representation of t**h**e four focus species was significantly greater in the expanded data set. The **p**roportion of pDNCs recovered for the main data set that re**m**ain valid and diagnostic in the ex**p**anded data set ranges from 13% In *C. miliaris* to 66% in *C. ebraeus*. This result confirms our expe**c**tations that the genetic diversity in the main data sets was clearly insufficient to propose a robust pDNCs-based diagnosis for species in Odonata and *Conus*.

The results of the iterated haplotype resampling of the four *Conus* species are shown on the figure 3. With the increasing captured genetic diversity of each species, the number of sampled independent pDNCs reduces only by 15-20%, and the increase of the proportion of EDV-pDNCs is almost linear, its local deviations from the linear regression closely paralleling those in the increasing PVS. The arbitrary confidence threshold of 0.8 is reached no earlier than when ∼50% of the haplotype diversity is sampled (*C. ebraeus, C. sanguinolentus),* and only when ∼66% and 74% of the haplotype diversity is sampled for *C*. *chaldaeus* and *C*. *miliaris* respectively.

**Figure 3.**
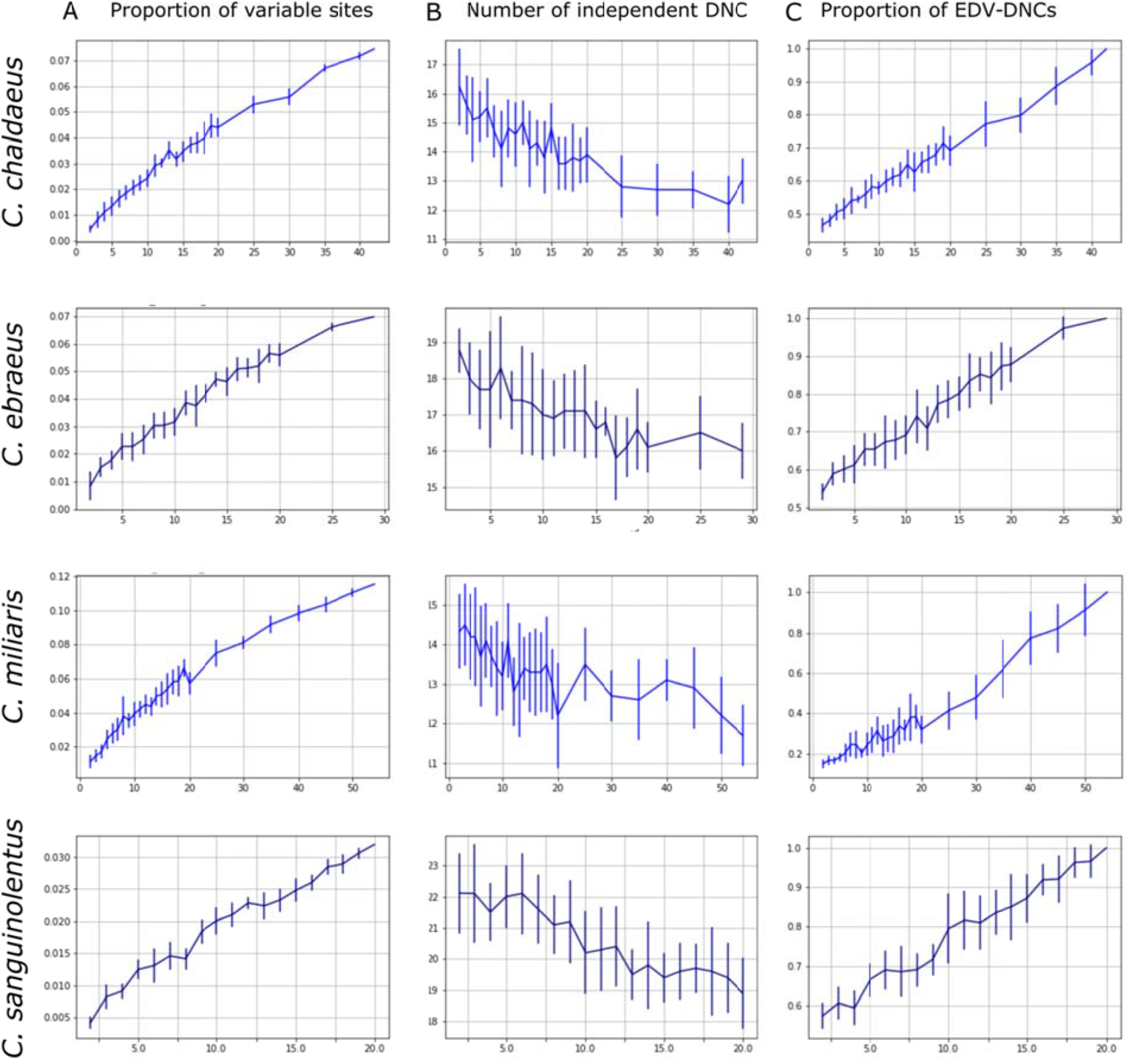
Haplotype resampling in four species of *Conus* and associated changes in **A**. Proportion of variable sites (PVS) in the DNA alignment; **B**. Number of independent pDNCs recovered, and **C**. proportion of EDV-pDNCs in the output. Vertical bars correspond to standard deviation calculated from 10 iterations of each sampling fraction.

#### Robustness of the pDNC-based diagnosis of supraspecific taxa

The column A on the figure 4 shows the changes of the genetic diversity (expressed through the PVS in the DNA alignment) depending on the sampled species fraction of Sclerurinae, *Synallaxis, Conasprella* and *Conus.* In the Sclerurinae the PVS continues to increase notably with each new species added to the data set until all 17 are sampled. In *Conus,* with only 25 species sampled (∼13% of the total number of species in the data set), ∼80% of the PVS in the COI alignment are recovered; however the subsequent increase of the PVS is consistent with linear, but not asymptotic, growth. In *Synallaxis* and *Conasprella,* the PVS seems to approach plateau, but only when 30 species (i.e. over 90% of the final species diversity in a data set) are sampled. The curves of increasing genetic diversity in the reduced data sets (with each species represented by single specimen) parallel those in the full data set. However, with the same number of species sampled, the PVS values in the reduced data sets were always lower.

**Figure 4.**
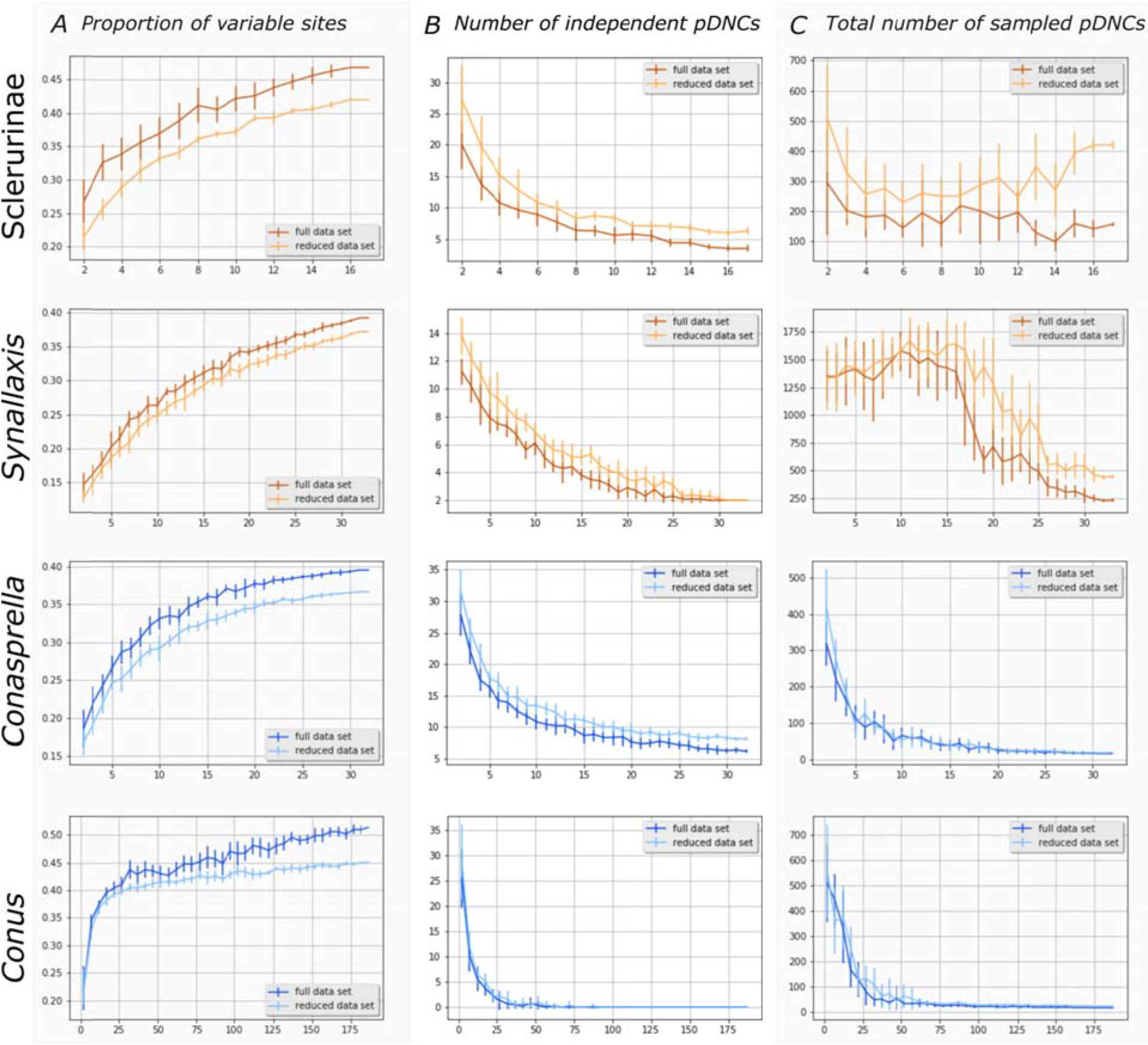
Species resampling in Sclerurinae, *Synallaxis, Conasprella* and *Conus* and changes associated with the sampling fraction in **A**. Proportion of variable sites (PVS) in the DNA alignment, **B**. Number of independent pDNCs and **C**. total number of pDNCs sampled. On each graph a darker line shows complete resampling regime (all sequences of the resampled species included in partial dataset), and a lighter line – reduced resampling (only one sequence per species included). Vertical bars correspond to standard deviation calculated from 10 iterations of each sampling fraction.

As the number of sampled species increases, both the mean number of independent pDNCs (figure 4, column B) and the mean standard deviation diminish, consistently with our expectations. However, only in *Conus* and *Synallaxis* the mean values reach a plateau, and the standard deviations reduce to zero. As expected, larger numbers of independent pDNC are recovered in the reduced data sets for Sclerurinae, *Synallaxis* and *Conasprella* for any sampling fraction, compared to the full data sets. Finally in both *Conasprella* and *Conus* (and in both reduced and full data set) the total number of recovered pDNCs decreases quickly when increasing the number of species sampled (figure 4, column C), and seem to closely reach a plateau. On the contrary, in both Sclerurinae and *Synallaxis,* no clear pattern in relation to the sampled fraction can be noted, except for larger number of pDNCs being recovered with a reduced sampling. This inconsistency suggests that the total number of recovered pDNCs is altogether a poor proxy of the molecular identity of a taxon.

We calculated which proportion of the pDNCs output for each sampling fraction were EDV-pDNCs (Figure 5). As expected, the initial proportion of EDV-pDNCs was negligible in all four analyses, with both the reduced and full resampling. In all analysed cases the proportion of the EDV-pDNCs was increasing very slowly. The threshold for the mean of 0.8 was reached at the species sampling fraction of ∼90% for Sclerurinae, *Synallaxis* and *Conasprella* and slightly earlier (at sampling fraction ∼50%) in *Conus;* however, these 50 % comprise as many as 93 species. The reduced sampling regime resulted in even lower proportion of the EDV-pDNCs: even when all species are sampled the proportion of EDV-pDNCs ranges from 0.14 (Sclerurinae) to 0.7 (*Conus*). Our results thus convincingly demonstrate that the multiple pDNCs proposed for the fraction of a data set generally do not hold when tested against the whole data set.

**Figure 5.**
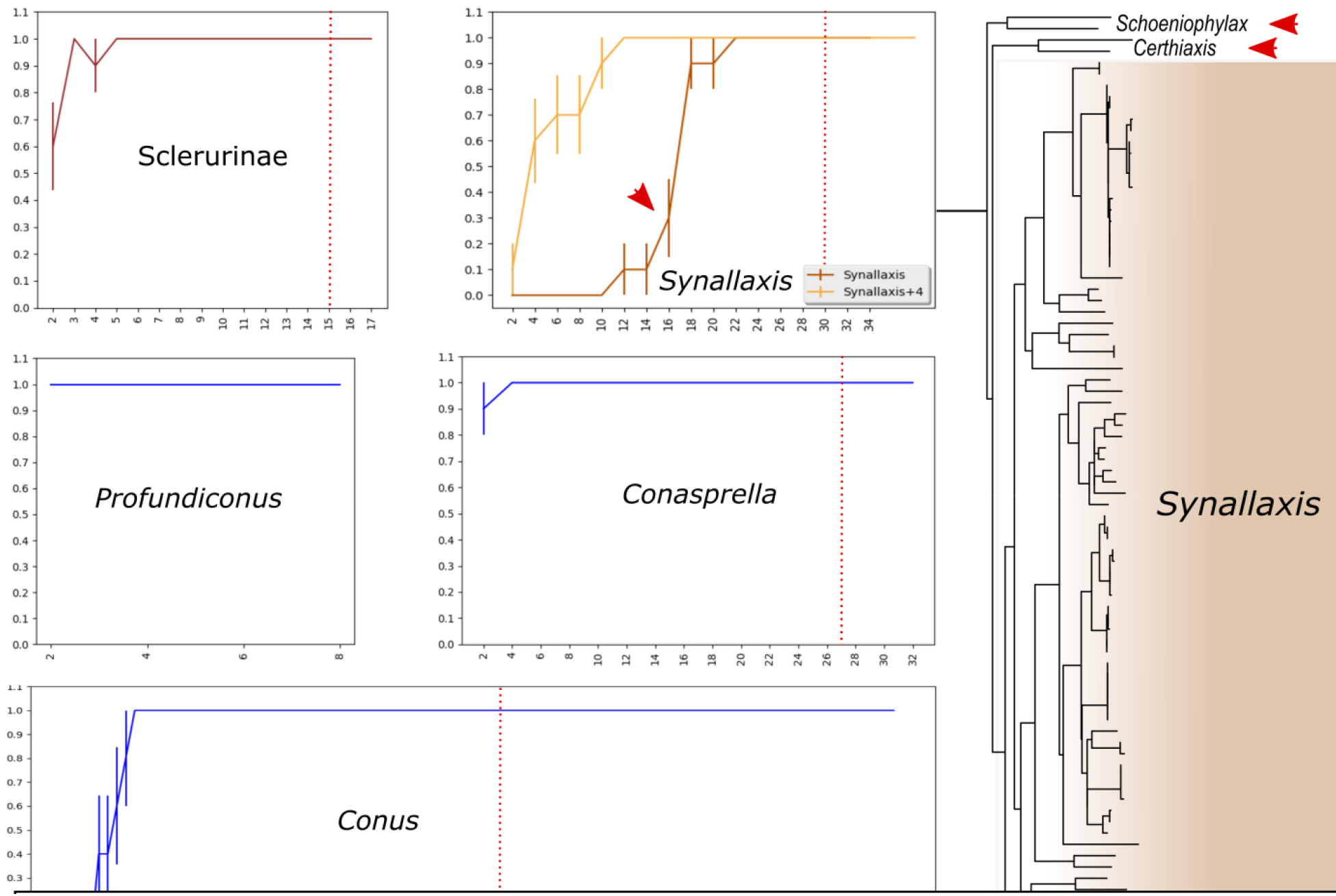
**A**. Species resampling and associated with it proportion of EDV-sDNCs in the output for Sclerurinae, *Synallaxis, Profundiconus, Conasprella* and *Conus.* Vertical bars correspond to standard deviation calculated from 10 iterations of each sampling fraction. Punctuated red line marks sampling fraction at which an arbitrary confidence threshold of 0.8 for pDNCs have been reached. **B**. ND1-based phylogenetic tree of *Synallaxis* and related Furnariidae taxa.

#### Diagnoses based on the sDNC for species-level taxa

The table 2 summarizes the sDNCs obtained for the 15 species of Odonata and 4 species of *Conus* based on the respective main data sets after validation on the artificially generated data sets. The sDNC-based diagnoses of 12 species of Odonata from the main data set (48 species) remained valid diagnostic combinations of these species in the scope of the much larger Odonata_4kCOI data set. In two of the remaining cases *(Crocothemis erythraea* and *Pseudagrion massaicum),* the sequences attribution to species appears to be inconsistent in the GenBank. The COI sequences of *Crocothemis erythraea* and *Crocothemis servilia* do not form reciprocally monophyletic groups due to the highly divergent sequence Jalala7 (Supplementary Fig. 2A), which was not included in the main Odonata data set. Similarly, the COI sequences of the *Pseudagrion massaicum* appeared intermixed, and in some cases identical to the sequences of *Pseudagrion tanganyicum* (Supplementary Fig. 2B), the latter also not included in the main Odonata data set. In the third case, the specimen RMNH-INS-508317 (also not included in the main data set) showed a much higher divergence from other sequenced specimens of *Trithemis stictica* (Supplementary Fig. 2C, sequences included in the main data set highlighted with violet). Although such value of genetic divergence does not contradict the assignation of this specimen to *T. stictica,* its addition to the data set notably increases the genetic diversity of the species, which does not fit the previously proposed diagnosis anymore. When the COI sequence of the RMNH-INS-508317 was added to the main data set and *Trithemis stictica* rediagnosed, the new sDNC (highlighted with green in the table 2) became a valid DNC for this species in the expanded data set as well.

**Table 2.** Secondary DNCs recovered for Odonata and Conidae species from main COI data sets.

| Species | sDNC | validity of<br>sDNC on<br>expanded<br>dataset |
| --- | --- | --- |
| *Pseudagrion bicoerulans* | [28:'G', 43:'C', 46:'C', 58:'C', 244:'G', 256:'T', 259:'C', 472:'G'] | Yes |
| *Anax ephippiger* | [40:'T', 94:'T', 142:'G', 349:'T', 370:'T', 454:'A', 481:'C', 502:'C', 521:'C'] | Yes |
| *Anax imperator* | [142:'G', 217:'T', 223:'C', 256:'T', 304:'C', 319:'T', 490:'T', 517:'G'] | Yes |
| *Anax speratus* | [79:'C', 139:'G', 145:'C', 178:'T', 244:'G', 319:'C'] | Yes |
| *Chlorocnemis abbotti* | [49:'G', 67:'C', 239:'T', 240:'C', 293:'A', 296:'A', 297:'A', 426:'G', 430:'A', 541:'C'] | Yes |
| *Crocothemis erythraea* | [85:'C', 220:'C', 268:'G', 277:'C', 310:'G', 331:'C', 427:'G', 502:'G'] | No |
| *Coryphagrion grandis* | [28:'T', 136:'G', 232:'T', 253:'T', 265:'C', 268:'C', 434:'C', 526:'G'] | Yes |
| *Crocothemis sanguinolenta* | [28:'T', 34:'T', 100:'G', 151:'G', 296:'C', 340:'C', 358:'T', 427:'G'] | Yes |
| *Orthetrum coerulescens* | [25:'C', 139:'C', 157:'G', 178:'T', 181:'G', 235:'C', 370:'G'] | Yes |
| *Platycypha caligata* | [29:'A', 30:'G', 40:'T', 182:'A', 214:'G', 233:'T', 397:'C', 403:'C', 410:'C', 439:'G'] | Yes |
| *Pseudagrion kersteni* | [100:'G', 112:'C', 157:'T', 223:'G', 232:'T', 235:'C', 403:'C'] | Yes |
| *Pseudagrion massaicum* | [7:'G', 19:'G', 37:'C', 151:'G', 187:'C', 400:'C', 484:'G', 505:'G'] | No |
| *Teinobasis alluaudi* | [7:'C', 34:'G', 154:'T', 217:'G', 340:'C', 343:'G', 448:'G', 457:'G'] | Yes |
| *Trithemis stictica* | [73:'C', 94:'T', 97:'C', 169:'G', 175:'T', 202:'G', 355:'T', 517:'G'] | No |
| ***Trithemis stictica*** | **[97:'C', 169:'G', 175:'T', 220:'C', 232:'T', 355:'T', 415:'T', 418:'G']** | **Yes** |
| *Orthetrum julia* | [13:'G', 76:'C', 124:'T', 175:'T', 227:'C', 277:'T', 361:'A', 487:'C', 505:'G', 514:'C'] | Yes |
| *Conus chaldaeus* | [35:'C', 94:'C', 97:'G', 118:'C', 127:'G', 289:'A', 586:'G', 637:'G'] | Yes |
| *Conus ebraeus* | [40:'C', 241:'G', 301:'C', 412:'C', 439:'G', 550:'T', 574:'G', 586:'A', 637:'A'] | Yes |
| *Conus miliaris* | [85:'G', 88:'A', 190:'T', 232:'G', 250:'C', 281:'C', 304:'G', 325:'G', 541:'G', 550:'T', 586:'G'] | Yes |
| *Conus sanguinolentus* | [40:'G', 121:'C', 307:'C', 346:'C', 445:'C', 565:'C', 571:'C'] | Yes |

Similarly, the sDNC-based diagnoses provided for the four species of *Conus* based on the main data set all appeared valid DNCs in the context of the expanded data set. The diagnoses retrieved for *Conus chaldaeus* based on only six unique COI haplotypes in the main data set remained valid in the context of the expanded data set including 41 unique COI haplotypes, in 20 independent MolD runs. Similar results were obtained also for *C. ebraeus* (12 unique COI haplotypes in the main data set VS 72 in the expanded one) and *C*. *miliaris* (21 VS 113 haplotypes). This result suggest that numerically, ∼20% of the haplotype diversity sampled (if expanded data set accepted as 100%) is enough to propose a reliable and reproducible DNA based diagnosis at the species level.

#### Diagnoses based on the sDNC for supraspecific taxa

The graphs of the changing proportions of EDV-sDNCs in the resampling outputs for Sclerurinae, *Synallaxis, Profundiconus, Conasprella* and *Conus* are shown on the Figure 5. The threshold of 0.8 is reached when 3 out of a total 17 species are sampled in Sclerurinae, 2 out of 32 for *Conasprella,* 14 out of 186 in *Conus* and 18 out of 32 in *Synallaxis* (brown line). Thus, in the four datasets this arbitrary reliability threshold is reached with less than 20% of species sampled, and almost all sDNCs retrieved for Sclerurinae, *Profundiconus, Conasprella* and *Conus* with a sampling fraction higher than 20% were EDV-sDNC. In the case of *Synallaxis,* however, the same threshold is reached only when 50% species are sampled. This genus also demonstrates the longest pDNCs among the analysed taxa (the shortest pDNC comprises 4 nucleotide positions – fig 2). The genus *Synallaxis* is the largest genus of the Furnariidae, nested within the crown group of the family, with a very short branch (Fig 5B). This topology of the tree suggests weak phylogenetic distinctiveness of *Synallaxis,* which explains the difficulties encountered in providing this genus a robust DNA-based diagnosis. In the treatment of Derryberry et al (2011), the genera *Schoeniophylax* and *Certhiaxis* are considered separate from *Synallaxis* (shown by red arrows of fig 5B); however, their status relative to *Synallaxis* is disputable (Claramunt 2014). If *Synallaxis* is considered in the expanded boundaries (i.e. including four species of the current genera *Schoeniophylax* and *Certhiaxis),* a more robust DNA-based diagnosis can be recovered, with the reliability threshold of 0.8 reached at ∼25% of the species sampled (orange line on the fig. 5A).

The sDNCs retrieved for the supra-specific taxa showed certain degree of inter-run variation (Supplementary table 9). Nevertheless, most positions remained conserved and shorter sDNCs comprised sub-sets of the nucleotide positions of the longer sDNCs output in other runs. Despite the variation of the length of the output sDNCs from run to run, because of the random effect associated with the sDNCs rating mechanism, their composition remained overall consistent, suggesting that they are closely comparable.

## Discussion

### MolD and other tools for diagnostic DNA characters recovery

The algorithm MolD is admittedly rough-and-ready – it does not take into account such parameters of a data set as the transition-transversion ratios or likelihoods of synonymous substitutions at the nucleotide positions included in the DNCs etc. These considerations are not within the scope of the present study, as it considers DNA-characters *as is* – i.e. not as evolving characters, but as fixed attributes of a set of specimens. We demonstrate that such model-free approach efficiently retrieves diagnostic combinations of nucleotides (DNCs) for pre-defined groups of DNA sequences in data sets of varying complexity. This algorithm is therefore scalable and versatile. Even for the taxa, where the shortest recovered pDNS comprised four nucleotide positions *(Synallaxis),* this pDNC was repeatedly resampled in the independent runs of MolD. This suggests that the default 10-20,000 recursions allow for exhaustive sampling of the informative nucleotide positions across the DNA alignment leading to reproducible pDNCs output. Subsequently, the tested algorithm to transform pDNCs into a sDNC allows compilation of a robust DNA-based diagnosis. All analyses described herein, including those on the extensive DNA data sets downloaded from GenBank, or those that involved taxonomic resampling, were carried out on one CPU of a moderately powerful laptop. Therefore, the program can easily run on virtually any decent computer with Python installed.

Most previous studies that report specific nucleotide substitutions have utilized one of the two softwares: function nucDiag in the SPIDER package for R (Brown et al. 2012), or CAOS (Characteristic Attribute Organization System – Sarkar et al. 2008 – Jörger and Schrödl 2013; Churchill et al. 2014; Zielske and Haase 2015, etc.). The former is only able to identify pure singlenucleotide diagnostic sites and can be efficient in the analysis of few species, but would generally fail when applied to data sets comprising hundreds of species, as pure single-nucleotide sites are less likely to exist as more and more sequences are included in the data set (Fedosov et al. 2019). CAOS (Sarkar et al. 2008) identifies private as well as pure characters of a focus taxon, and also allows compilation of composite DNA characters. This functionality makes CAOS a powerful and versatile program to identify DNA barcodes that can be directly used in DNA based descriptions (Goldstein and DeSalle 2011; Jörger and Schrödl 2013; 2014). However, CAOS implements a tree based approach, which, on one hand, has an obvious merit – the identified characters, (termed Characteristic Attributes or CAs) are meaningful in evolutionary context, but, on the other hand, also has a drawback – only clades in the input tree can be diagnosed. This requirement in practice may not be met (Goldstein & DeSalle 2011): the new taxa may not be monophyletic when the tree is reconstructed based on a single molecular marker (the one used to establish the diagnosis). It has been shown that up to 25% of the species are not monophyletic in the analyses of monolocus data sets (Funk & Omland 2003), and this proportion is expected to be even higher for more genetically heterogeneous higher rank taxa.

MolD retrieves pure and composite nucleotide characters (similar to CAOS), but implements an alternative approach and does not postulate that the predefined groups are monophyletic for the DNA fragment used for the diagnosis. Therefore, the monophyly of these groups must have been previously established and our goal is not to verify the validity of these groups (whatever the criteria used to delimit them), but to propose a DNA-based diagnosis for each of them. Same diagnosis should ideally be shared by all members of a diagnosed lineage. To achieve that we abstained from using private characters (i.e. nucleotide positions polymorphic among the sequences of a focus taxon). However, we make use of pure and nested characters. We find that avoiding the use of composite characters in the DNA-based diagnoses as suggested by Jörger and Schrödl (2013) is conceptually poorly justified, when dealing with already delineated taxa. When monophyly (i.e. shared ancestry) of a diagnosed taxon has been explicitly demonstrated, all characters shared by the taxon’s members (and only such characters are considered herein) should be considered as descending from a common ancestor (for detailed justification see Davis & Nixon 1992). Equally, we disagree with the statement that only or preferentially synapomorphic characters should make their way into diagnoses. The purpose of a diagnosis is to communicate identity of a taxon, and in line with it, a diagnosis focuses on character states, but not on their evolutionary history. To our knowledge, most, if not all, traditional diagnoses comprise *informative* morphological characters irrespective of their apomorphic or plesiomorphic nature, or being homologous or analogous. A diagnosis in general, and *a fortiori* a tree-independent diagnosis, could thus be compared to identification key – following the key enables allocation of a specimen to certain taxon, but the consecutive dichotomies of a key are not expected to match the events in the taxon’s evolutionary history.

### Taxonomic sampling and robustness of the DNA-based diagnoses

The major impediment to the proposition of molecular diagnoses on a regular basis is (the concern of) the lack of robustness of the diagnoses, because of their sampling-dependent nature. Theoretically, in order to claim that a diagnostic character of some taxon is truly fixed, every single individual of a this taxon would need to be examined, which apparently will never be feasible (Wiens & Servedio 2000). Tripp and Lendemer (2014) suggested that preferably 10 vouchers of any new taxa should be sequenced along with at least 15 closest relatives, which should enable ‘sampling broad enough to encompass the phylogenetic, geographical and ecological diversity within each taxon’ (Tripp & Lendemer 2014: 970). The numbers provided by Tripp & Lendemer (2014) look somewhat arbitrary, and clearly imply restricted taxonomic scope of the analysis.

Our results show that reliable diagnoses can be recovered for species in data sets covering taxa as diversified and ancient as Odonata. Remarkably, out of 15 Odonata species diagnoses compiled with only 5% of the final species diversity covered (main data set VS expanded data set) 12 remained valid in the context of the expanded data set. Only in the case of *Trithemis stictica,* the COI-based diagnosis recovered for the main Odonata data set was not valid against the expanded data set, despite all sequences assignments were correct. In this sole case, a notably divergent haplotype of the focus taxon was missing in the main data set. Thus, it is not the number of species or sequences included in the data set, but rather proper representation of the extremes of the diversity range that can be seen as a main requirement for the taxonomic sampling leading to proposition of a reliable molecular diagnosis. Essentially, same requirement defines sufficient dataset for the species delimitation based on monolocus data, or DNA barcoding. This requirement cannot be unequivocally translated into number of haplotypes and species, because some taxa are more difficult to diagnose than others. Nevertheless, when the effect of the haplotype sampling was tested, and the species coverage was the same (main data set VS expanded data set), we were able to repeatedly compile valid diagnoses for the four *Conus* species based only on <20% of their final haplotype diversity. As well, species coverage of ∼20% was usually sufficient to repeatedly compile robust DNA based diagnoses for the analysed supraspecific taxa. However, as many as 50% of the final species number needed to be resampled to reliably diagnose *Synallaxis* as currently conceived.

We demonstrate that the reliability of the DNA-based diagnoses can be estimated using simple informatics toolkit – mainly thanks to the formalized and universal language of the DNA. The traditional morphological diagnoses, even theoretically, cannot be challenged in a similar manner, because they are mainly based on the difficult to formalize taxon-specific features and are also subject to researcher bias (Fujita et al. 2012). In this prospect, DNA-based diagnoses compiled following a standardized protocol in a thoughtfully designed data set should be a more reliable descriptor of a taxons’ identity, compared to the traditional morphological diagnoses. In any case, revision of a morphological diagnosis is a common practice, when novel data become available. Similarly, the DNA based diagnosis would in most cases remain a reflection of state of the art in understanding molecular identity of a taxon.

### pDNCs versus sDNCs

The results of taxonomic resampling demonstrate convincingly that pDNCs can hardly be useful for diagnosing taxa, unless the taxonomic coverage is close to or exceeds 90 %, and each species is represented by multiple DNA sequences. Unless these very strict requirements are met, the diagnoses would not be sufficiently robust even in a context of known taxonomic diversity of a taxon. On the contrary, the required level of confidence is reached with notably lower sampling fraction with sDNCs. We have demonstrated that the developed step-wise diagnosis validation algorithm is capable to efficiently model the genetic diversity missing in the analysed data set, thus allowing for robust DNA-based diagnoses to be retrieved even with partial taxon sampling. We also confirm the conclusions of Rach et al. (2008) and Bergmann et al. (2009; 2013) that single genetic marker is sufficient to confidently diagnose species, and in most cases, supra-specific taxa, and the taxon’s identity can be conveyed by a relatively short sDNC. The criteria of robustness used in the present study for sDNCs (a score of 100 or 85 in three consecutive validation steps for species or supra-specific taxa respectively) were chosen arbitrarily. Such parameters may be too strict, and sufficient robustness may be achieved even with more relaxed criteria, and correspondingly, with shorter sDNCs.

While the algorithm that we used to compile sDNCs in the present study is novel, the nature of the sDNC is essentially the same as of a barcode in character-based DNA barcoding (DeSalle et al. 2005; Rach et al. 2008): a set of informative nucleotides in the alignment is compiled to convey a molecular identity of a taxon. However, the common practice of the character-based DNA-barcoding is to provide barcodes involving the same set of nucleotide positions across all taxa in a data set, so the barcodes also have same length. It can be best illustrated on the studies of Odonata, for which the species barcodes comprised 23 (ND1 – Rach et al. 2008) or 29 nucleotide positions (both COI and ND1 – Bergmann et al. 2013). On the contrary the approach implemented herein outputs a unique set of nucleotides for each analysed taxon, so the sDNCs vary in length. In the case of Odonata species sDNC are notably shorter compared to the barcodes of Rach et al. (2008) and Bermann et al. (2013) – on average 6.77 and 7.66 nucleotide positions for ND1 and COI, respectively. Beside this technical discrepancy, there are differences in the purpose of the diagnostic combinations of nucleotides: in the DNA barcoding studies they serve for identification of a query specimen, whereas in a framework of taxonomic study they aim at providing a better formal diagnosis for a taxon. Therefore, the methodology of the character-based DNA barcoding requires a tool to retrieve barcodes to be complemented by another tool which matches query sequences to a set of barcodes for taxonomic identification. No such tool is required when the purpose of the diagnostic combination of nucleotides is the taxonomic description. However, as the nature of DNA barcodes and sDNCs is the same, these two applications of the DNCs are perfectly compatible: diagnoses provided in the taxonomic description can be readily used for specimen identifications by means of character-based DNA barcoding. We consider this operational compatibility with adjacent fields of DNA-based systematics as a main strength of the developed algorithm, providing a better integrity of the DNA-based systematics and taxonomic studies.

### What if no DNCs retrieved?

The failure to recover a robust diagnosis may be due to a technical issue (incorrect attribution of a sequence to a taxon, inclusion of a contaminated sequence to a dataset, or a misalignment), or have a biological reason. It is essential that the DNA alignment used as an input for MolD is checked carefully to ensure that the sequences are genuine, naming is consistent throughout the data set, and the alignment is accurate. It is possible that no diagnosis can be returned for the highly diversified and thus heterogeneous taxa and/or for those that have diverged too little from the closest relative. The former case can be exemplified in our data by the subfamily Furnariinae, for which MolD was not able to recover sDNC, and no pDNCs were identified in the extended data set. Conversely, the example of a pitfall at the species level is the species pair *Pseudagrion acaciae* – *P. nyloticum* in the Odonata data set: the COI sequences of these species have also been misidentified by BOLD classifier (Bergmann et al. 2013). The failure in both these cases can be explained by the fact that the resolution limit of the respective genetic marker has been reached (see Hickerson et al. 2006). Similar failures are more likely in the taxa, where standard DNA barcode markers are known to evolve slowly (e.g. COI in sponges and cnidarians – Erpenbeck et al., 2006), or to produce gene trees highly incongruent with species trees (Funk & Omland 2003; Mutanen et al. 2016). In such cases, alternative genetic marker(s) should be analysed. Alternatively, single DNA-based diagnosis can be provided for the weakly differentiated species pair, such as *Pseudagrion acaciae* – *P. nyloticum.* The extremely low degree of divergence between these species suggests that their delimitation and status should be revisited with the use of specific approaches (such as the multilocus species delimitation – Pante et al. 2015a) that proved efficient when incomplete lineage sorting is anticipated.

### How a DNA-based diagnosis should be presented

In the current study we address the methodological impediments which hamper use of the DNA sequence data in the taxonomic descriptions. As the earlier recommendations on the incorporation of DNA-sequence data in taxonomic practice (e. g. Jörger & Schrödl 2013; Tripp & Lendemer 2014) did not address these aspects, we here elaborate on them specifically.

#### Characters in DNA based diagnosis

In the present analysis we take for granted that a summary of specific nucleotides at the specific positions of DNA alignment is a most usable manner of DNA data presentation. We demonstrate that a redundant set of such informative positions shared by all members of a focus taxon (a sDNC) allows for more robust diagnoses, and should be preferred over a summary of independent diagnostic characters (a pDNC-based diagnosis). The genetic marker and primers used to generate DNA sequences should be stated explicitly in the subsection of Materials and Methods dedicated to the taxonomy (i.e. separately from the phylogenetic analysis). The reported nucleotide positions are relative to the forward primer used for PCR; therefore, to avoid inconsistency, standard primers (such as the Folmer’s barcode primers for COI) should be preferred. In the case that alternative primer combinations are used, the obtained positions should be recalculated to match either the standard ‘0’ position defined by a respective commonly used forward primer, or the specific index in the coordinates of complete mitochondrial genome. We show that the design of the data set, namely representation of both the focus taxon and non-focus taxa, is essential for retrieving a robust DNA-based diagnosis. Therefore, we suggest that 1) a brief characterization of the analysed data set should be included in the sub-sections of the Materials and Methods dealing with taxonomy, 2) the number of sequenced individuals (for species), or number of sequenced individuals and the number of species represented (for supra-specific taxa) should be stated explicitly in the diagnosis. Additionally, the nucleotide alignment used to generate DNA-based diagnoses should be provided in supplementary data.

#### Algorithm

We developed the algorithm MolD specifically to use DNA data in taxonomic descriptions of taxa. It has some advantages over CAOS, and only outputs DNA-based diagnosis if it meets pre-defined criteria of robustness. Nevertheless, the sets of characteristic attributes generated by the p-gnome module of CAOS (Sarkar et al. 2008), such as the barcodes presented by Rach et al. (2008) or Bergmann et al. (2013) can also be used, in case no private characters are included. Although, ready to use as is, MolD implementation in the stand-alone software is anticipated, aiming primarily for stable, user-friendly and platform-independent release. In future we also envision developing a web-based version of MolD, to be integrated in the toolkit of DNA barcode data bases, such as BOLD.

### MolD within the workflow of a contemporary systematics study

The crucial operational assumption behind the proposed approach is that the taxa to be diagnosed have already been delineated in the integrative taxonomy framework. This notion aligns with the methodological review by Goldstein and DeSalle (2011) who advocated for the necessarily independent procedures of the taxon discovery and taxonomic establishment. Therefore, while inability to provide a concise DNA-based diagnosis to a taxon may point at technical error at the taxa delimitation performed upstream, it is not and should not be a criterion for reassessing taxa delimitation any further than correcting an error. Therefore, the use of the DNA characters in taxonomic descriptions will not erase taxonomic impediment, as the taxonomic expertise will still be required for integrative taxa delimitation. Nevertheless, the use of DNA in formal descriptions would relieve taxonomists and save time at the stage of preparing descriptions, while improving their quality.

What Goldstein and DeSalle (2011) have identified as a future challenge is ‘to reconcile the precise mechanics of barcoding analysis with the empirical and philosophical rigor of systematics’. In this context, the algorithm implemented in MolD is capable of enhancing transition from discovery of a new or redefinition of known lineages to their formal description. The research workflow when MolD is expected to be most efficient, is when a phylogenetic analysis supported by the morphological, ecological and geographic data has led to redefinition of the scope of taxa, and the subsequent taxonomic revision of the (re)defined taxa is to be accomplished. The methodology of the coupled discovery and subsequent taxonomic description of taxa is becoming increasingly widely used and encouraged (Tripp & Landemer 2014), and we envision the developed algorithm for compilation of DNA-based diagnoses to become a useful tool in taxonomic studies.

## Data availability

All the alignments, output files and scripts used in the present study are submitted to Github (https://github.com/SashaFedosov/MolD) and will be made publicly available upon acceptance of the paper.

## Acknowledgments

Authors are grateful to Dr Heike Hadrys (ITZ, Hannover) for providing the COI and ND1 data sets on the order Odonata.

## Supplementary materials

Data available from https://github.com/SashaFedosov/MolD. The legend to the provided scripts, input and output files is provided in the github.com repository as well.

**Supplementary figure 1**. Length distribution of the pDNCs in Odonata and *Conus* species.

**Supplementary figure 2**. RAxML phylogenetic trees (COI) of *Crocothemis erythraea* and *C*. *servilia* (**A**), *Pseudagrion massaicum* and *P. tanganyicum* (**B**) and *<u>Trithemis stictica</u>* (**C**). COI sequences included in the main Odonata_COI data set highlighted with violet. Sequences, precluding compilation of a sDNC-based diagnosis valid for the expanded data set shown with red arrow.

**Supplementary table 1**. Metrics of the DNC based diagnoses for Odonata species (ND1, 316 positions).

**Supplementary table 2**. Metrics of the DNC based diagnoses for Odonata species (COI, 541 positions).

**Supplementary table 3**. Metrics of the DNC based diagnoses for Odonata families (data sets COI and ND1).

**Supplementary table4**. Metrics of the DNC based diagnoses for *Conus* species (COI, 658 positions).

**Supplementary table 5**. Metrics of the DNC based diagnoses for *Conus* subgenera (CO1, 658 positions).

**Supplementary table 6**. Metrics of the DNC based diagnoses for Conidae genera (CO1, 658 positions).

**Supplementary table 7**. Taxon-shared VS informative nucleotide positions in three codon positions.

**Supplementary table 8**. Validity of the pDNCs recovered for Odonata and Conidae species from main COI data sets on the respective expanded data sets.

**Supplementary table 9**. Inter-run variation of the sDNCs for taxa in analysed data sets.

